# Glucocorticoids Induce the Expansion of an Immature Human CFU-E Population

**DOI:** 10.1101/722850

**Authors:** Ryan Ashley, Hongxia Yan, Nan Wang, John Hale, Brian M. Dulmovits, Julien Papoin, Meagan Olive, Namrata D. Udeshi, Steven A. Carr, Adrianna Vlachos, Jeffrey M. Lipton, Lydie Da Costa, Christopher Hillyer, Sandrina Kinet, Naomi Taylor, Narla Mohandas, Anupama Narla, Lionel Blanc

## Abstract

Despite effective clinical use, the mechanistic bases for the regulation of human erythropoiesis by glucocorticoids remain poorly understood. Here, we employed an erythroid culture system to differentiate primary human CD34^+^ cells isolated from control peripheral blood, cord blood and patients with Diamond Blackfan anemia (DBA), as model of an erythropoietic disorder treated with glucocorticoids. We found that the response to Dexamethasone is dependent on the developmental origin of the source of CD34^+^ cells, specifically increasing the expansion of CD34^+^ cells from peripheral blood but not cord blood. We also demonstrated that Dex treatment leads to the expansion of a novel immature colony-forming unit-erythroid (CFU-E) population in peripheral blood which is uniquely responsible for the proliferative effects observed during human adult erythropoiesis. Dex treatment of peripheral blood CFU-E cells also induced p57^Kip2^ expression while patients with DBA resistant to steroids had altered p57^Kip2^ expression that did not increase with Dex. Furthermore, shRNA knockdown of p57^Kip2^ reduced proliferation and abrogated the effects of Dex. Finally, proteomics of Dex-treated CFU-E from peripheral blood and cord blood additionally revealed novel Dex targets in the erythroid system. Altogether, these results provide novel insights into the regulation of human erythropoiesis by glucocorticoids.

**Graphical Abstract:** 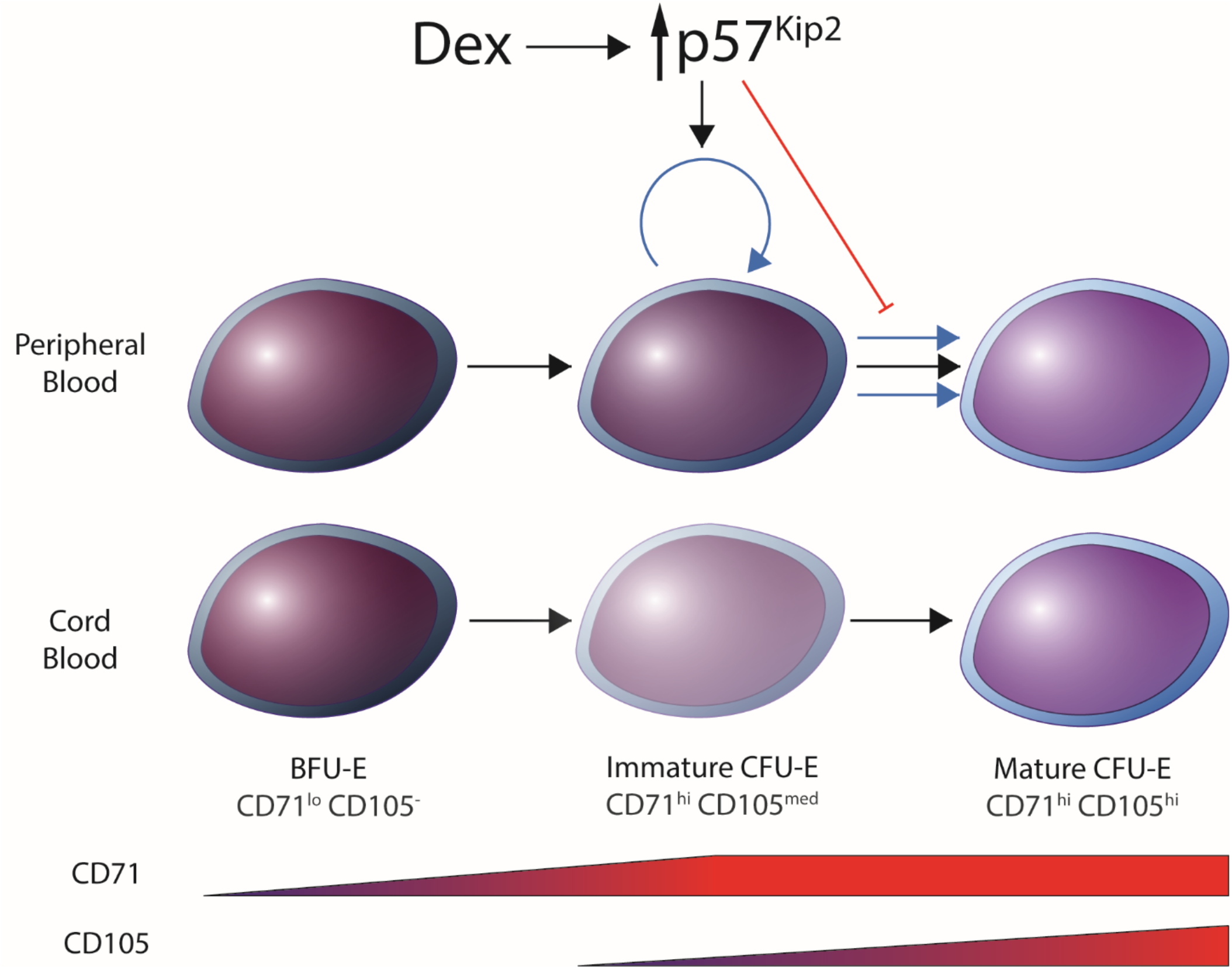

## Introduction

Glucocorticoids such as dexamethasone (Dex) are well known clinically to promote erythropoiesis and used as therapeutic agents in the treatment of disorders of red blood cell production including Diamond Blackfan anemia (DBA) (1–3). These molecules interact with the glucocorticoid receptor (GR) allowing for the nuclear translocation of the resulting complex which binds DNA at glucocorticoid response elements (GREs) and ultimately activates gene transcription (4).

DBA is an inherited bone marrow failure syndrome with an incidence of ~5 cases/million live births, characterized by red cell aplasia, a range of physical anomalies, developmental bone defects and cancer predisposition (5, 6). More than 70% of the patients diagnosed with DBA have defects in ribosome biogenesis due to mutations in genes encoding ribosomal proteins. In addition, mutations in the *GATA1* transcription factor, a key regulator of erythroid development, and *TSR2*, a pre-rRNA processing protein, have recently been identified in several DBA families (7, 8). The genetic landscape of DBA is heterogeneous but genotype/phenotype correlations have been noted in association with mutations in the *RPL5* and *RPL11* ribosomal protein in patients (9).

The standard of care for patients with DBA after the first year of age is glucocorticoids. Notably, a majority of treated patients have an increase in red cell production and exhibit reduced dependency on blood transfusion. However, the therapeutic dosage is extremely variable between patients and many patients become refractory to treatment over time. Once patients are glucocorticoid-refractory they become dependent on chronic red blood cell transfusions unless they enter remission or undergo a curative hematopoietic stem cell transplant (2).

The actions of glucocorticoids have been well studied in many disease contexts, but their specific mechanisms of action in the erythroid system in both healthy individuals and patients with DBA still remains to be fully elucidated. Several studies have demonstrated that glucocorticoids act at the erythroid progenitor level but the precise stages of erythroid differentiation at which they exert their effects is not yet known (10–13). This is in part due to the significant heterogeneity of erythroid progenitor populations and the different markers and model systems that are used for studying erythropoiesis and the effects of glucocorticoids.

Significant progress has been made in our understanding of terminal erythroid differentiation, from the proerythroblast to the reticulocyte stages, but a major limitation to our understanding of human red cell production has been the lack of a detailed cellular and molecular characterization of the early erythroid progenitors, specifically burst-forming unit-erythroid (BFU-E) and colony-forming unit-erythroid (CFU-E) cells. Classically, these populations have been identified on the basis of their ability to form erythroid colonies in semisolid medium colony forming assays (CFAs) and more recently, they have been identified immuno-phenotypically by the evaluation of surface markers with flow cytometry (14–18). Furthermore, progenitor cells derived from patients with DBA have been shown to uniquely contribute to defective erythropoiesis (19).

A further difficulty in evaluating the impact of glucocorticoids on BFU-E and CFU-E progenitors is due to important differences in murine versus human erythroid differentiation. In mice, the vast majority of research has focused on fetal liver progenitors, with a detailed characterization of early and late populations of fetal liver BFU-Es (20–22). Indeed, several studies performed on murine fetal liver cells have suggested that Dex acts at the BFU-E stage (10, 11, 13, 20). However, a recent study in mice has reported that Dex enhances the maintenance of proliferative CFU-E by upregulating p57^Kip2^, a member of the Cip/Kip cyclin-dependent kinase inhibitor protein family. Under these conditions of slowed S phase progression, the proliferative CFU-E population is maintained and there is a delayed differentiation to the less proliferative proerythroblast (Pro-EB) stage. However, it is not clear as to how fetal murine erythroid progenitors translate to humans, with regards to both the potential *ex vivo* heterogeneity amongst BFU-E and CFU-E populations and more importantly, the presence of different erythroid progenitor subpopulations in human bone marrow. These limitations have made it difficult to determine the stage of erythroid progenitor development at which glucocorticoids are acting in patients with DBA.

In the present study, we focused on the role of Dex during normal and disordered human erythropoiesis. The expansion of CD34^+^ hematopoietic stem and progenitor cells (HSPCs) isolated from adult peripheral blood (PB) were markedly increased following Dex treatment but surprisingly there was no increase in the proliferation of CD34^+^ HSPCs derived from cord blood (CB). Using highly enriched populations of BFU-E and CFU-E, we found that the proliferation of PB-derived, but not CB-derived, CFU-E was markedly increased by Dex. However, given the significant heterogeneity in human erythroid progenitors isolated on the basis of commonly used surface markers, we evaluated new markers and found that the combination of CD34, CD36, CD71 and CD105 allowed for a more precise discrimination of BFU-E and CFU-E subpopulations. We identified a transitional PB CFU-E progenitor population (CD34^+^CD36^+^CD71^hi^CD105^med^) as Dex-responsive but did not detect responsiveness in BFU-E, either in PB or CB. The differential Dex-mediated proliferative effect on PB CFU-E was associated with an upregulation of p57^Kip2^ in these cells and not in CB-derived CFU-E. Notably, we found that CFU-E isolated from patients with steroid-resistant DBA do not upregulate p57^Kip2^ in response to Dex. Finally, using mass spectrometry-based quantitative proteomic analysis, we identified the NR4A1 transcription factor as a novel regulator of the responsiveness to Dex in adult human CFU-E and that it was not upregulated in CB CFU-E.

Taken together, we have identified a novel population of immature CFU-E in adult PB that is Dex-responsive and reveal that the upregulation of p57^Kip2^ and NR4A1 is critical for this response. Furthermore, we find that the glucocorticoid refractory anemia in patients with DBA is associated with a lack of p57^Kip2^ upregulation. These findings will open new avenues for the development of specific therapeutic strategies for these patients.

## Results

### Dex increases the proliferation of adult human CFU-Es

In erythroid cultures of HSPCs from healthy adults and from patients with DBA it has previously been shown that Dex increases proliferation of erythroid cells (23, 24). However, the differentiation stage at which Dex exerts its effects is still unclear. Using a serum-free expansion media that allows for effective erythroid differentiation of HSPCs under steady state conditions without requiring Dex, we studied the effects of Dex on human erythropoiesis (18, 25). CD34^+^ cells isolated from adult PB or from CB were cultured in the presence of 100nM of Dex. There is a 7-fold increase in the proliferation of CD34^+^ cells from adult PB following Dex treatment (**Figure 1A**). In marked contrast and very surprisingly we noted that there is no increase in the proliferation of CD34^+^ cells derived from CB. In fact, we observed a decrease in proliferation implying marked developmental differences in the response of adult and cord blood CD34+ cells to Dex (**Figure 1A**).

**Figure 1:**
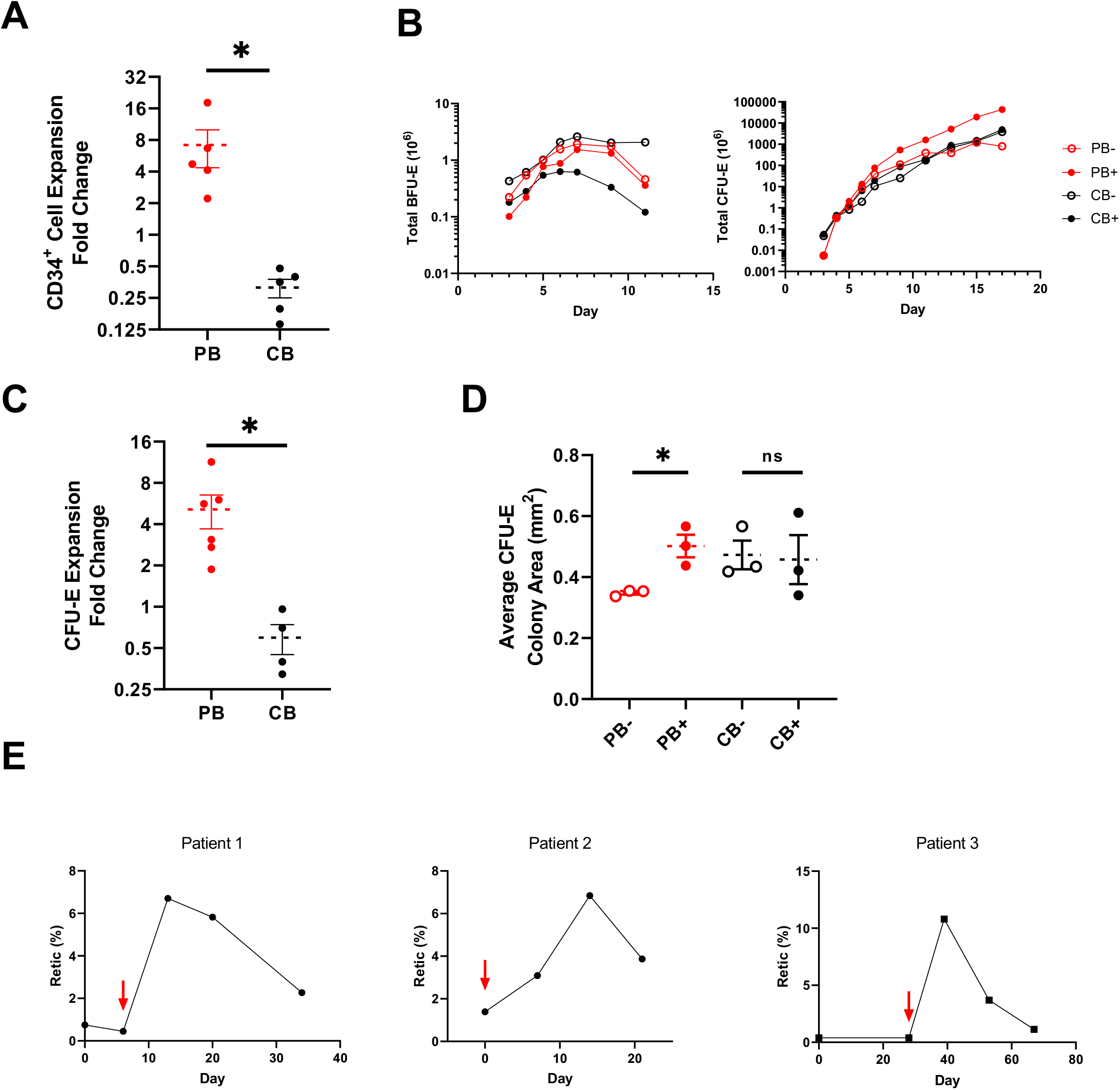
Dex enhances the proliferation of human peripheral blood CFU-Es. (A) Fold change in total expansion of PB and CB (n = 5) derived CD34^+^ cells upon expansion with Dex vs. untreated control. (B) Total numbers of BFU-E and CFU-E from PB and CB cultures during expansion with or without Dex. (C) Fold change in total expansion of purified CFU-Es derived from PB (n = 6) and CB (n = 4) upon expansion with Dex vs. untreated control. (D) Average area of colonies formed from purified CFU-Es derived from untreated PB and CB and Dex treated PB and CB (n = 3). (E) Retic response of DBA patients treated with prednisone. Treatment was initiated on days marked with red arrows and dose was titrated down on following time points. PB is shown in red while CB is shown in black. Dex treated cultures are shown as solid while untreated cultures as shown as open or hatched. Data for A, C, and D are shown as mean ± SE. Data shown in B is from 1 of 3 independent experiments performed. *P < 0.05, by 2-tailed Student’s *t* test (A, C and D).

We then attempted to further elucidate the erythroid differentiation stage at which Dex acts by using our recently developed experimental strategy for the characterization of the erythroid progenitors, BFU-E and CFU-E, based on the surface expression levels of GPA, IL3R, CD34, and CD36. The absolute number of BFU-E and CFU-E generated from 10^6^ CD34^+^ cells from both PB and CB were enumerated on the basis of cell surface marker expression in the absence and presence of Dex in culture (**Figure 1B**). Starting at day 7, the number of BFU-E (defined as GPA^-^ IL3R^-^ CD34^+^ CD36^-^) cells decreases for both PB and CB CD34^+^ cells in the absence of Dex and the extent of this decrease is enhanced in the presence of Dex. Interestingly, the decrease in number of BFU-E is more pronounced for CD34^+^ cells from CB. In marked contrast, the absolute number of CFU-E (defined as GPA^-^ IL3R^-^ CD34^-^ CD36^+^) increases over time and this increase is more pronounced for CD34^+^ cells originating from adult peripheral blood in the presence of Dex. Interestingly, the percentage of GPA^+^ cells representing the terminally differentiating erythroblasts decreases in the presence of Dex implying a delayed transition of the progenitor cells to terminal erythroid differentiation (**Supplemental Figure 1A**). Terminal erythroid differentiation is also delayed at D11 of culture in PB-treated cells, as assessed by flow cytometry using the surface markers a4-integrin and Band 3 as well as by reduced hemoglobinization **(Supplemental Figure 1B)**.

To further validate that the CFU-E is the population primarily responding to Dex, we sorted CFU-E from peripheral and cord blood cultures of CD34^+^ cells and studied their proliferative potential in serum-free expansion media. As shown in **Figure 1C**, the proliferative capacity of purified CFU-E from adult blood increases 5-fold in the presence of Dex compared to control cultures without Dex. Once again, Dex has little to no effect on the proliferation of purified CFU-E from cord blood, supporting the idea that the response to Dex is linked to the the specific developmental stage of erythroid progenitors. We performed the same experiments using increasing concentrations of Dex on sorted populations of erythroid progenitors derived from adult PB cultures and observed that the effect on proliferation is maximal at 100nM (**Supplemental Figure 2**).

Functional assays using methylcellulose cultures with Epo only revealed that treatment with Dex markedly increases the colony size of PB CFU-E but has very little or no effect on CB CFU-E colony size **(Figure 1D, Supplemental Figure 3)**, in spite of the fact that in the absence of Dex the colony size of purified CB-derived CFU-E was larger than that generated by PB CFU-E. Taken together, these data imply that only human CFU-E from adult PB respond to Dex by increasing proliferation.

Previous studies have showed that *in vivo* it takes at least two weeks *in vivo* for the normal bone marrow to produce reticulocytes from the BFU-E stage and about 7 to 10 days from the CFU-E stage (26–28). We hypothesized that if CFU-E is the progenitor population that responds to glucocorticoids *in vivo*, then patients treated with steroids should present with reticulocytosis in less than two weeks. We followed patients with DBA over a one-month period before and after treatment with prednisone. We observed that in each patient the reticulocyte count increases within seven to eleven days after initiation of the treatment strongly suggesting that *in vivo*, the CFU-E is indeed the population responsive to glucocorticoids **(Figure 1E-F)**. We further noticed a decline in the reticulocyte response in a similar time frame, in association with a decrease in the dose of prednisone administered to the patient.

### Dex targets a subpopulation of adult CFU-Es

We previously described that based on surface expression of CD34 and CD36 we can obtain highly enriched population of BFU-E (GPA^-^ IL3R^-^ CD34^+^ CD36^-^) and CFU-E (GPA^-^ IL3R^-^ CD34^-^ CD36^+^) (18). More recently we reported a transitional progenitor population defined as GPA^-^ IL3R^-^ CD34^+^ CD36^+^ which is more predominant during differentiation of adult PB than that of CB (29). Notably, the kinetics of progression through these differentiation states were also altered with Dex treatment **(Figure 2A)**.

**Figure 2:**
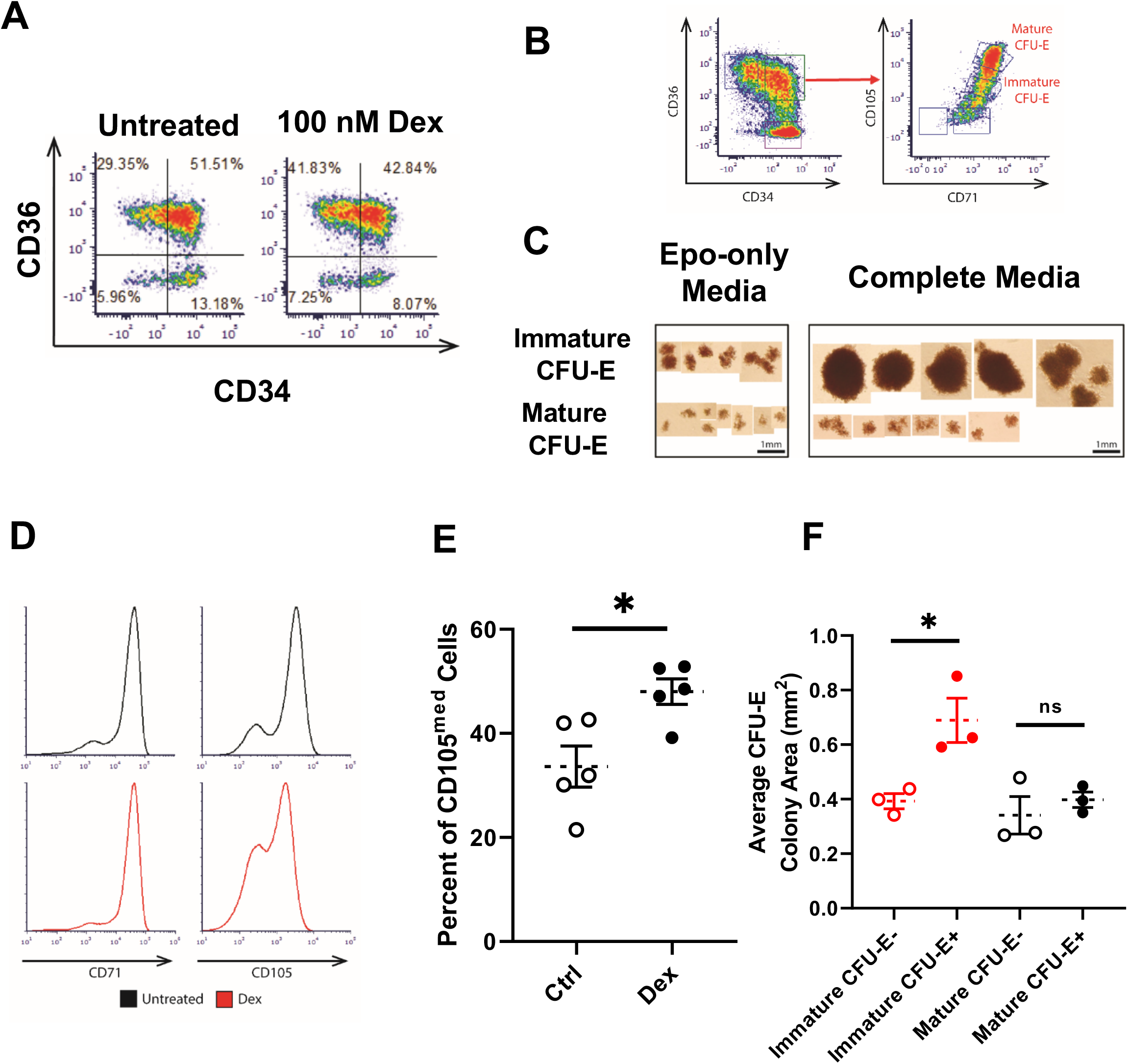
Dex targets a transitional subpopulation of peripheral blood CFU-Es. (A) Representative flow cytograms of PB derived CD34^+^ cells treated with or without Dex at day 4 of expansion by CD36 and CD34 expression. (B) Gating strategy to define transitional erythroid progenitor populations. (C) Representative images of colonies formed by cells purified from populations after growth in Epo-alone or complete medium CFA. (D) Representative flow cytograms of PB-derived CD34^+^ cells treated with or without Dex at day 4 of expansion by CD105 expression. (E) Quantification of CD105^hi^ cells in cultures of PB-derived CD34^+^ cells treated with or without Dex at day 4 of expansion. (F) Area of colonies formed from purified populations in Epo-alone media CFA with Dex treatment (n = 3) or without Dex treatment (n = 3). Immature CFU-E are shown in red and mature CFU-E are shown in black. Dex treated cultures are shown as solid while untreated cultures as shown as hatched. Data for A, B, C and D represent 1 of 3 independent experiments. Data for E and F are shown as mean ± SE. *P < 0.05, by 2-tailed Student’s *t* test (E and F).

Having identified two phenotypically different CFU-E populations (CD34^+^ CD36^+^ and CD34^-^ CD36^+^) we sought to determine if we can identify additional cell surface markers that would provide further insights into the heterogeneity of these CFU-E cell populations. We focused on CD71, the transferrin receptor, and CD105, endoglin, which both demonstrate significant differences in their RNA expression during erythroid differentiation at the progenitor stages **(Supplemental Figure 4)**.

Based on the expression patterns of CD71 and CD105 on CD34^+^ CD36^+^ cells we identified two distinct populations – CD71^hi^ CD105^med^ and CD71^hi^ CD105^hi^ **(Figure 2B)**. We then sorted these different cell populations and performed colony forming assays in the presence of either Epo alone, to produce colonies with the traditional definition of CFU-E, or in complete medium with SCF, Epo, IL3, IL6, G-CSF and GM-CSF. Interestingly, while both sorted cell populations from CD34^+^ CD36^+^ generate colonies in the presence of Epo only **(Figure 2C, left panel)**, only CD71^hi^ CD105^med^ show marked responsiveness to SCF in complete medium, while SCF has little or no effect on CD71^hi^ CD105^hi^ cells **(Figure 2C, right panel)**. CD71^hi^ CD105^hi^ cells from CD34^-^ CD36^+^ populations behave similarly to those from CD34^+^ CD36^+^ populations. Based on these findings we propose that CD71^hi^ CD105^med^ be termed as immature CFU-E and CD71^hi^ CD105^hi^ as mature CFU-E. When treated with Dex, PB-derived CD34^+^ cells preferentially maintain this immature CFU-E population. Indeed, while both untreated and treated cells express comparable levels of CD71 on their surface, ~ 50% of the PB cells treated with Dex are still CD105^med^ in comparison to untreated controls, ~ 30% of which are CD105^med^ at day 4. **(Figure 2D-E)**. Importantly, immature CFU-E cells functionally respond to Dex by increasing their colony size in methylcellulose culture system **(Figure 2F)**. In marked contrast, mature CFU-E respond very marginally to Dex in the same functional colony-forming assays and this increase is not statistically significant. Taken together, these data demonstrate that in human erythropoiesis, Dex treatment preferentially maintains the immature CFU-E progenitor population for an extended period of time to increase its proliferative capacity.

### Dex induces negative cell cycle regulation

Based on our improved immuno-phenotyping methodology, we studied populations of BFU-E, immature CFU-E and mature CFU-E derived from human PB. After staining these populations with Hoechst 33342 for cell cycle analysis, we observed that PB-derived immature CFU-E treated with Dex specifically show a decreased percentage in the S phase population that is not observed in the BFU-E or the mature CFU-E after treatment **(Figure 3A, Supplemental Figure 5)**.

**Figure 3:**
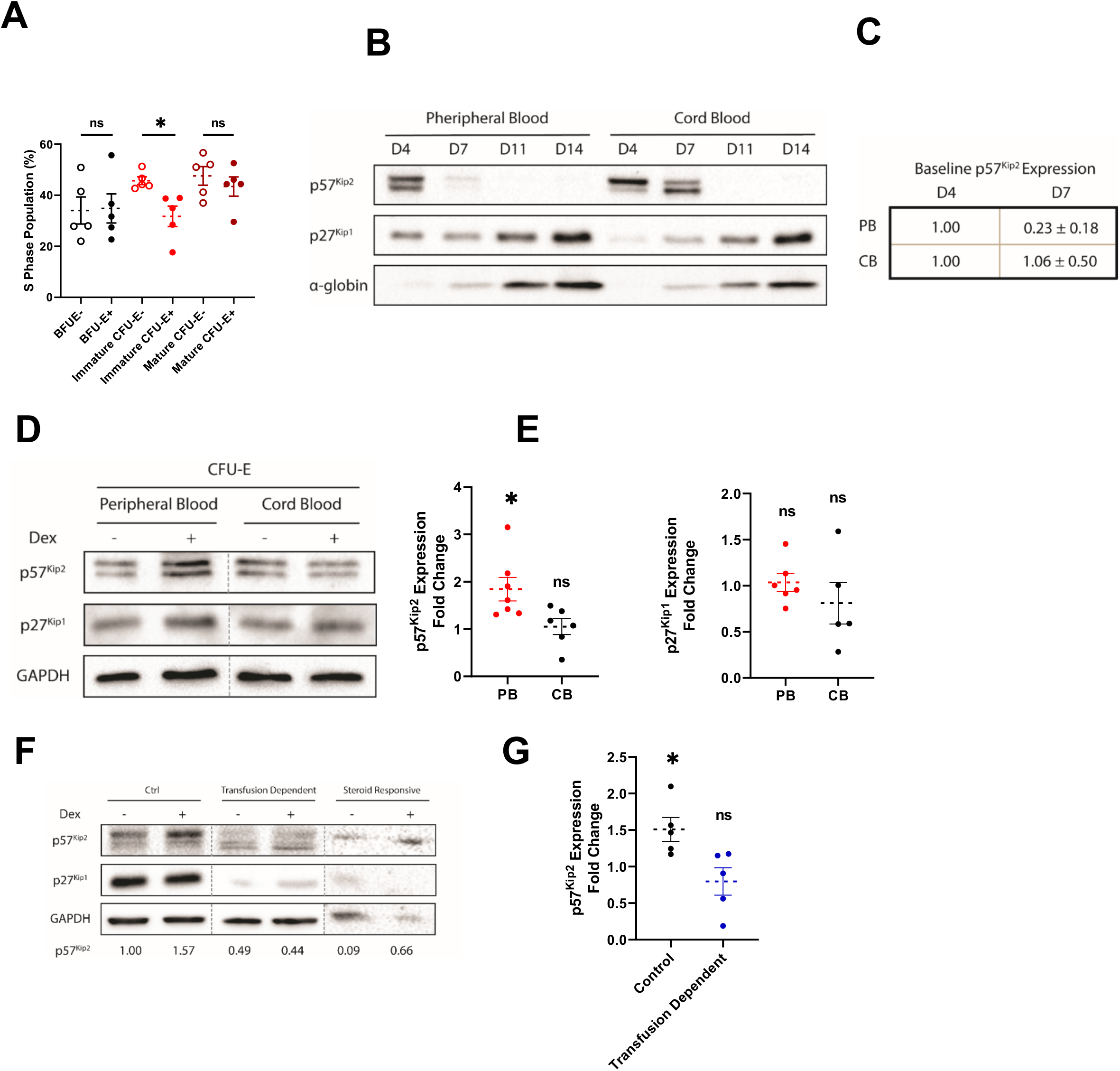
p57^Kip2^ expression is increased with Dex and dysregulated in DBA. (A) Quantification of the population of cells in S phase for BFU-E, immature CFU-E and mature CFU-E derived from PB treated with or without Dex and stained with Hoechst 33342 (n = 5). (B) Expression of p57^Kip2^ and p27^Kip1^ over the time course of differentiation. (C) Quantification of differences in baseline expression of p57^Kip2^ between day 4 and day 7 of culture for PB and CB (n = 5) (D) Expression of p57^Kip2^ and p27^Kip1^ by purified CFU-Es derived from PB and CB treated with or without Dex. (E) Fold change of p57^Kip2^ expression in purified CFU-Es derived from PB (n = 7) and CB (n = 6) and p27^Kip1^ expression in purified CFU-Es derived from PB (n = 6) and CB (n = 5) treated with Dex vs. untreated control. PB is shown in red while CB is shown in black. (F) Expression of p57^Kip2^ and p27^Kip1^ by erythroid progenitors derived from DBA patients and healthy controls. Expression of p57^Kip2^ relative to GAPDH is quantified below. (G) Fold change of p57^Kip2^ expression in CD34^+^ cells derived from transfusion dependent DBA patients or PB from healthy controls treated with Dex vs. untreated control (n = 5). Data for B, D and F represent 1 of 3 independent experiments. Data for A, C, E and G are shown as mean ± SE. *P < 0.05, by 2-tailed Student’s *t* test (A, C, E and G).

### p57^Kip2^ expression is increased by Dex in human CFU-Es

We then investigated whether this change in cell cycle status by Dex is due to increased expression of p57^Kip2^ as was recently described for murine erythroid progenitors (10). We first evaluated the expression levels of p57^Kip2^ protein during erythroid differentiation of CD34^+^ cells derived from PB and from CB **(Figure 3B)**. We noted that p57^Kip2^ is expressed in the erythroid progenitors and then its levels decrease during terminal stages of erythroid differentiation (days 7 to 14). Interestingly the kinetics of its expression pattern is different depending on the origin of the CD34^+^ cells **(Figure 3B)**. While p57^Kip2^ levels are reduced by about 80% at day 7 of differentiation in PB-derived CD34^+^ cells, its expression persists in cells derived from CB **(Figure 3B, C)**. Conversely, the expression of p27^Kip1^, another member of the Cip/Kip family is minimal in progenitor cells and increases dramatically in terminally differentiating erythroblasts **(Figure 3A)**. RNA sequencing data of enriched populations of erythroid cells at different differentiating stages are consistent with the noted protein expression patterns **(Supplemental Figure 4)**. This is also consistent with previous work that implicated p57^Kip2^ in maintaining quiescence at the stem and progenitor cell stages, while p27^Kip1^ has other functions during terminal differentiation (30, 31).

Based on these findings, we treated sorted CFU-E cells with Dex for 24 hours and evaluated the expression levels of p57^Kip2^ in these cells. In PB CFU-E treated with Dex we observed increased expression of p57^Kip2^ while CB CFU-E showed no changes in expression level following Dex treatment **(Figure 3D, E)**. The levels of p27^Kip1^, on the other hand, are minimally affected by Dex in CFU-Es from either of the two sources **(Figure 3D, E)**. Altogether, these results suggest that p57^Kip2^ is a target of Dex in adult human CFU-Es, as it has been shown in mouse CFU-Es (10).

### p57^Kip2^ expression is altered in erythroid progenitors from transfusion-dependent patients with DBA

We further hypothesized that the resistance to glucocorticoids in patients with DBA could be mediated, at least in part, by p57^Kip2^. To test this hypothesis, we isolated CD34^+^ cells from the peripheral blood of normal controls and transfusion-dependent patients with DBA. We observed that while the levels of p57^Kip2^ are increased in the blood from normal controls after treatment with Dex, there is a minimal change in its expression levels in patients with DBA **(Figure 3F, G)**. As expected, the levels of p27^Kip1^ are unchanged in the presence or absence of Dex, although they are significantly decreased in DBA samples with defective differentiation. In patients that are responsive to steroids, however, we could notice an upregulation of the expression levels of p57^Kip2^ after treatment of the HSPCs with Dex **(Figure 3F)**. Taken together, these results suggest that p57^Kip2^ associated cell cycle changes play a role in steroid resistance during human erythropoiesis.

### Genetic ablation of *CDKN1C* (p57^Kip2^) confers Dex insensitivity

In order to test whether p57^Kip2^ is necessary for the observed effects of Dex on erythroid progenitors, we employed lentiviral shRNA constructs targeted against p57^Kip2^ mRNA **(Figure 4A)**. Following transduction with a luciferase targeting shRNA control construct, peripheral blood-derived CD34^+^ cells retain their Dex sensitivity and display significant increases in cell growth, as we observed in cultures without lentiviral transduction **(Figure 4B)**. However, in cells transduced with p57^Kip2^ targeting shRNA constructs, there is a severe deficiency in proliferation that is not rescued following treatment with Dex **(Figure 4B)**. We also performed similar experiments to knockdown p27^Kip1^ and did not notice a significant effect of the knockdown on progenitor cells, only minor changes in terminal differentiation, consistent with our understanding on the relative roles of p27^Kip1^ and p57^Kip2^ during normal human erythroid differentiation **(Supplemental Figure 7A, B)**. Interestingly, we noted that while genetic depletion of p57^Kip2^ led to marked decrease in cell proliferation, it hastened terminal erythroid differentiation as assessed by GPA expression levels at day 14 of culture, independent of Dex treatment **(Figure 4C)**. Altogether, these data indicate that p57^Kip2^ is critical in controlling the balance between proliferation and differentiation during human erythropoiesis.

**Figure 4:**
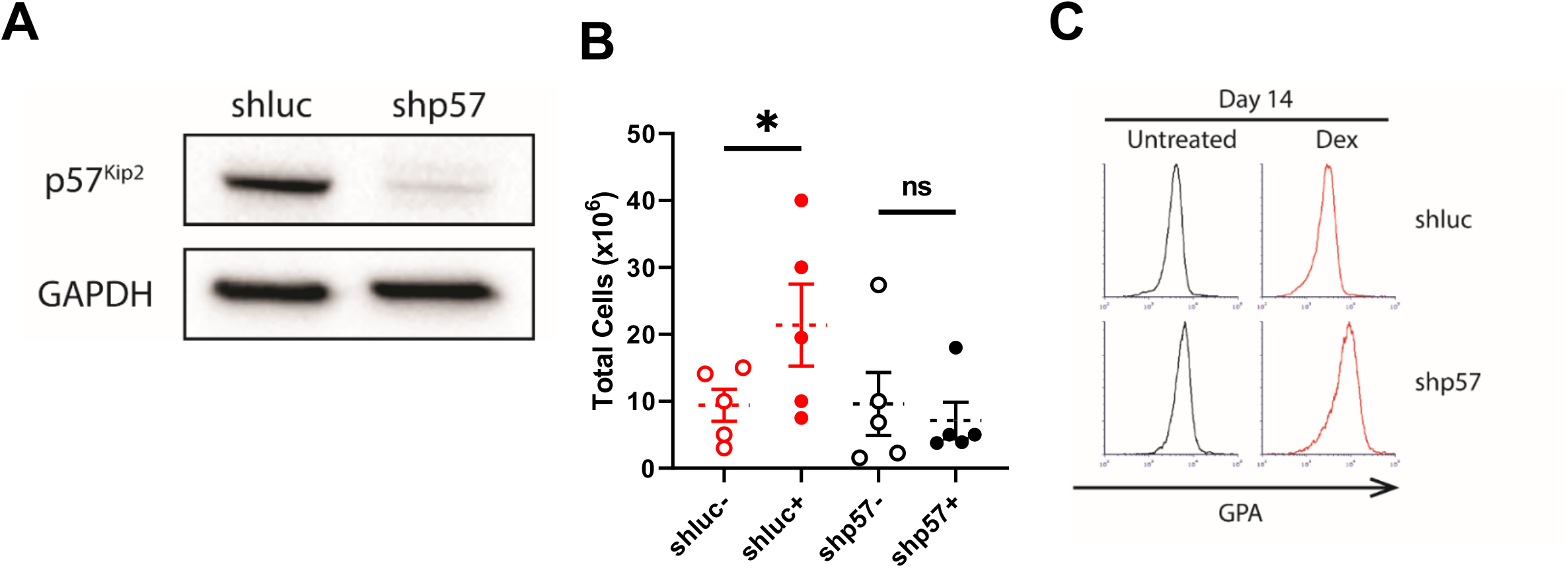
Genetic ablation of *CDKN1C* (p57^Kip2^) confers Dex insensitivity and accelerates differentiation. (A) Expression of p57^Kip2^ in PB transduced with shRNA lentiviral constructs targeting luciferase and p57^Kip2^. (B) Total cell proliferation of PB CD34^+^ cells transduced with lentivirus for shRNA knockdown of p57^Kip2^ and luciferase control with or without Dex treatment (n = 5). (C) Representative flow cytograms of PB CD34^+^ cells transduced with lentivirus for shRNA knockdown of p57^Kip2^ and luciferase control with or without Dex treatment at day 14 of differentiation. shRNA against luciferase cultures are shown in red and shRNA against p57^Kip2^ cultures are shown in black. Dex treated cultures are shown as solid while untreated cultures as shown as open. Data for A and C represent 1 of 3 independent experiments. Data for B are shown as mean ± SE. *P < 0.05, by 2-tailed Student’s *t* test (B).

### Proteomic studies highlight NR4A1 as novel Dex target

Having demonstrated that p57^Kip2^ plays a critical role in the Dex response observed in human adult CFU-Es, we explored if other proteins could be involved in the mechanism of action of glucocorticoids using a proteomics approach. After 5 days of *in vitro* culture, we sorted PB- and CB-derived CFU-E and treated them with or without 100nM Dex for 24 hours. These cells were processed for proteomic analysis by digestion with trypsin and labeling of resulting peptides with isobaric tandem mass tag reagents (TMT10) for relative quantification. Labeled peptides were subsequently analyzed by liquid chromatography-tandem mass spectrometry. Proteomics quantified 10,045 proteins in PB samples and 10,028 number of proteins in CB samples **(Supplemental Table 1)**. Both PB and CB samples showed regulation of several proteins including PER1, which is responsible for the repression of glucocorticoid-induced transcriptional activity (**Figure 5A, B**) (32). However, CFU-E derived from peripheral blood demonstrated specific upregulation of several proteins following Dex treatment. Among these is NR4A1 (Nuclear Receptor Subfamily 4 Group A Member 1), a negative cell cycle regulator (33) which is noted to be among the top 20 proteins upregulated in CFU-E derived from PB following Dex treatment **(Figure 5C)** but not in CB **(Figure 5C)**. Additionally, both the glucocorticoid receptor and p27^Kip1^ were identified in both PB and CB samples but were not significantly regulated by Dex, as expected from our expression data **(Supplemental Figure 4)**. p57^Kip2^ was not identified with this approach, likely due to its low abundance in late progenitors.

**Figure 5:**
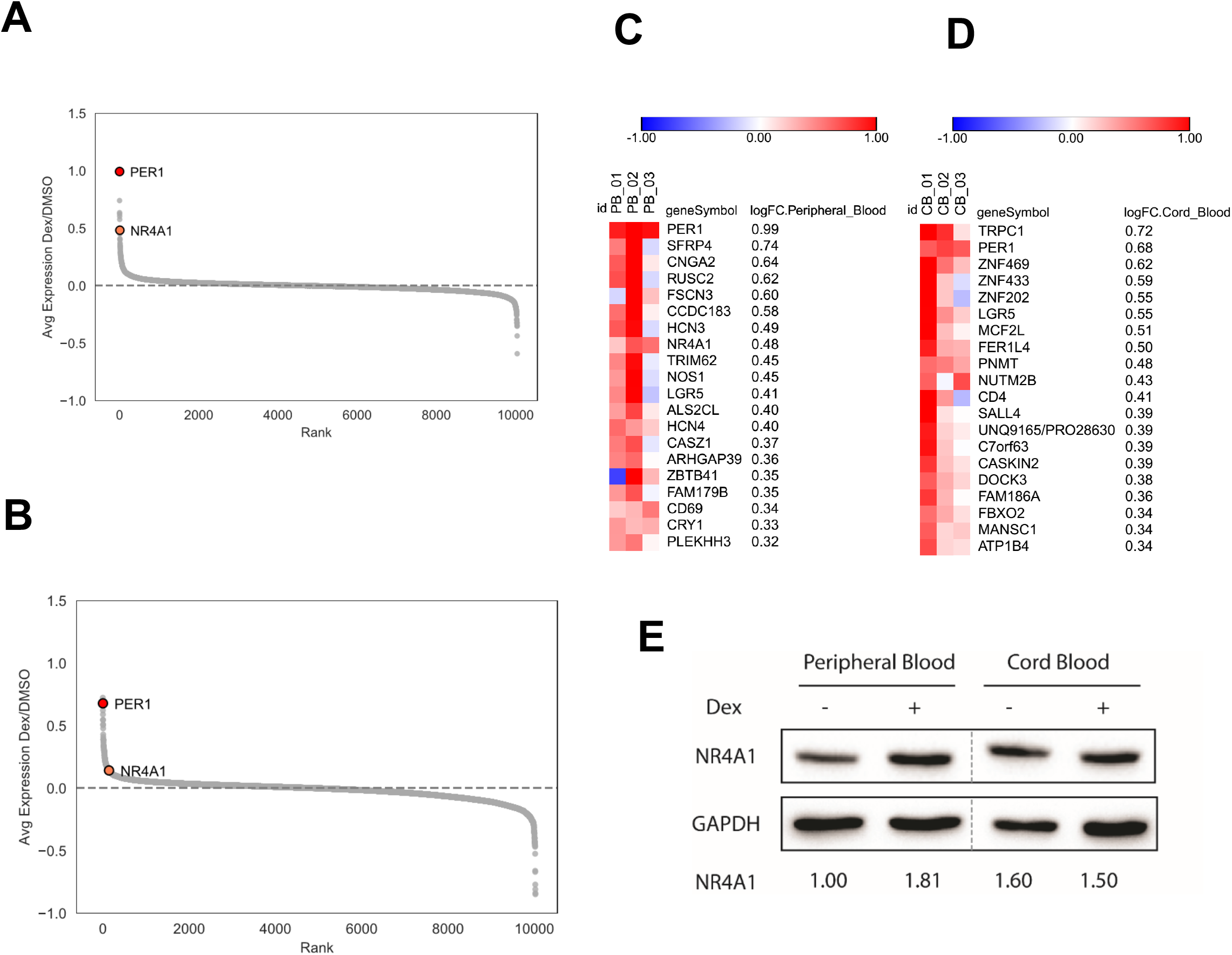
Proteomic studies highlight NR4A1 as novel Dex target in peripheral blood. (A) Ranked average fold change plots of differences in protein expression between Dex treated PB. (B) Ranked average fold change plots of differences in protein expression between Dex treated CB. (C) Top 20 upregulated proteins in Dex treated PB. (D) (C) Top 20 upregulated proteins in Dex treated CB. (E) NR4A1 expression in purified CFU-Es derived from PB and CB. Expression of NR4A1 relative to GAPDH is quantified below. Data for E represent 1 of 3 independent experiments.

To gain further insights into the role played by NR4A1 in the response to Dex, we evaluated its expression levels during erythropoiesis following Dex treatment. We observed that the levels of NR4A1 increase for human PB-derived erythroid cells, but not in CB-derived erythroid cells (Figure 5E). These data suggest that NR4A1 may be an additional important downstream regulator in the erythroid response to Dex.

## Discussion

The present study focused on defining the role of Dex during human erythropoiesis has generated several novel insights. The finding that Dex markedly enhanced the erythroid proliferation of CD34^+^ cells from adult PB but not the proliferation of CD34^+^ cells derived from CB was unexpected and very surprising. In conjunction with our previous finding of different trajectories in the transition from BFU-E to CFU-E between CD34^+^ cells from adult PB and CB this finding implies that Dex has differential effects on transitional erythroid progenitor populations that is dependent on the developmental state (29). The use of enriched populations of human BFU-E and CFU-E cells enabled us to define that the CFU-E derived from adult human PB and not BFU-E are responsive to Dex. This finding is not consistent with the long-held view based on extensive studies on murine fetal liver derived erythroid cells that BFU-E population is Dex responsive. It should be noted, however, that findings from a recent murine study has also implicated CFU-E to be Dex responsive (10). The present findings raise the important issue regarding attention that needs to be paid to the developmental origin of hematopoietic stem and progenitor cells in addition to the culture methods used in the study of factors that regulate human erythropoiesis. Interestingly, the identification of human CFU-E as target of Dex is consistent with the time course of the reticulocyte response seen *in vivo* in patients with DBA following treatment with prednisone.

The marked heterogeneity of human erythroid progenitors has long been recognized (34, 35). Indeed, it has been known for decades that erythroid progenitors give rise to colonies of different sizes and morphology in the methylcellulose culture system which led to the concept of large, intermediate and small BFU-E colonies and that the small BFU-E forming cells further differentiate into CFU-E (36). The recent advent of single cell technologies has also enabled finer analysis of these varied populations in terms of both steroid sensitivity and cell fate decisions (12, 37).

We have previously shown that human BFU-E and CFU-E can be immuno-phenotypically defined based on the surface expression of IL3R, GPA, CD34 and CD36. However, this characterization did not allow us to fully resolve the well documented heterogeneity of human erythroid progenitors. In the present study, we document that in addition to these markers, CD71 and CD105 can be used to further immuno-phenotypically discriminate sub-populations of erythroid progenitor cells. Using this method, we identified several new transitional progenitor populations with distinct phenotypes, including two distinct populations of CFU-Es, designated immature and mature CFU-Es.

Exploring the effects of Dex on human erythropoiesis using this optimized immunophenotyping assay, we found that treatment of adult cell populations resulted in a specific early increase and maintenance of the immature CFU-E population. Importantly, this specific population was found to be highly responsive to Dex both in terms of proliferation in CFAs and induction of a decrease percentage of cells in the S phase of the cell cycle. This cell cycle phenotype was due to an increase in p57^Kip2^ expression by Dex which we found was essential for the effect of Dex on proliferation. This was demonstrated by a large decrease in proliferation combined with accelerated differentiation after shRNA knockdown of p57^Kip2^. This demonstrates a clear link between cell cycle progression and differentiation of erythroid progenitors, which aligns with recent work that identifies the role of cell cycle status in cell fate decisions (38). Importantly, we also found that patients with DBA that are transfusion-dependent present altered levels of p57^kip2^, and these levels do not change after treatment. One caveat in this experiment relies on the fact the levels of p57^Kip2^ were not assessed over time. The other limitation is the number of patients with DBA that are steroid-responsive who are accessible for studies, since these patients do not need chronic blood transfusions. However, we observed a trend towards elevated levels of p57^Kip2^ following Dex treatment *in vitro*, serving as proof of concept for the importance of this cell cycle regulator in the response to steroids.

We propose that through expansion and maintenance of the immature CFU-E population, Dex generates larger numbers of mature CFU-E. It is critical that this increase in proliferation occurs before entering the Pro-EB stage since large increases in proliferation cannot take place due to the more tightly linked progression of differentiation and cell division during terminal erythroid differentiation. Additionally, our proteomics data revealed novel Dex targets that may offer additional insight into how Dex influences the cell cycle and increases erythroid proliferation. Of these targets, many have been shown to be involved in cell cycle regulation including PER1 and NR4A1, indicating that these proteins may also be involved in inducing negative cell cycle regulation alongside p57^Kip2^ (32, 39). It should also be noted that NR4A1 is involved in TGF-β signaling, which is also aberrant in DBA, and could be involved in corticosteroid resistance mechanisms (40). Importantly, we also documented that Dex had no effect on human BFU-E prior to further maturation.

The present findings have direct relevance for the clinical use of corticosteroids for the treatment of hypo-proliferative anemias, most notably DBA. While corticosteroid use stably increases red cell mass and induces remission in many patients with DBA, others see little effect or cannot continue with glucocorticoids due to the side effects of treatment. We anticipate that our new insights into the mechanism of action of glucocorticoids on human erythropoiesis will allow for the development of new targeted drugs and treatment strategies to induce a sustained erythroid effect in more patients with fewer side effects. In summary, we have introduced a new method for defining human erythroid progenitor populations with distinct phenotypic differences and found that dexamethasone predominately acts at a specific transitional stage between late BFU-E and mature CFU-E stages. These findings contribute to both our further understanding of erythroid progenitor biology and potential new treatment strategies for erythroid disorders.

## Materials and Methods

### Isolation and culture of CD34^+^ cells

PBMCs were separated using Lymphoprep (Stem Cell Technologies) and CD34^+^ cells were purified with anti-CD34 microbeads and MACS Columns (Miltenyi Biotec) using manufacturer protocol. CD34^+^ cells were cultured at a density of 10^5^ cells/mL in a serum free expansion media as previously described (25), with a base media of StemSpan SFEM (Stem Cell Technologies) initially supplemented with 100 ng/mL stem cell factor (SCF), 10 ng/mL interleukin 3 (IL3), 0.5 U/mL erythropoietin (Epo), 4 μL/mL lipid mixture 1 (Sigma Aldrich), 2 mM L-glutamine and 200 μg/mL transferrin. Beginning at day 7, the dose of Epo was increased to 3U/mL and at day 11 the dose of transferrin was increased to 1 mg/mL. Dexamethasone (Sigma) was added to cultures as indicated at 100 nM, its dose of maximal effect as supported by data in **Supplemental Figure 2**.

### Flow cytometry and cell sorting

Erythroid progenitors were examined using surface markers as previously described (18). Erythroid progenitors were analyzed at day 4 of differentiation where 10^5^ cells were stained with an antibody cocktail containing anti-IL3R PE-Cy7, anti-glycophorin A (GPA) PE, anti-CD34 FITC, anti-CD36 APC, anti-CD71 Alexa Fluor 700, and anti-CD105 Brilliant Violet 421 (BD Biosciences) for 15 minutes at room temperature. BFU-E were defined as GPA^-^ IL3R^-^CD34^+^ CD36^-^, CFU-E as GPA^-^ IL3R^-^ CD34^-^ CD36^+^, and transitional progenitors as GPA^-^ IL3R^-^ CD34^+^CD36^+^. Dead cells were excluded from further analysis with 7-aminoactinomycin D (BD Biosciences) staining. Analysis was preformed using a BD Fortessa flow cytometer and FCS Express 6 and FlowJo 10. BFU-E, CFU-E and transitional progenitors were sorted for downstream experiments using a BD FACSAria at a low pressure with a 100 μm nozzle.

### Colony forming assays

Sorted erythroid progenitors were seeded to methylcellulose media H4230 (Stem Cell Technologies) supplemented with Epo alone at 0.5 U/mL or complete methylcellulose media H4435 (Stem Cell Technologies) at 200 cells/mL. BFU-E and CFU-E colonies were counted and measured at day 14 and day 7 of culture respectively. Area of colonies was determined by modeling each colony as an ellipse and measuring its major axis **a** and minor axis **b** to calculate area by the formula **A = πab/4**. Dex was added to cultures as indicated at 100 nM.

### Cell cycle staining

For live cell cycle staining for examining cell cycle profile, erythroid progenitors were incubated with 5 mg/mL Hoechst 33342 for 3 hours at 37°C prior to staining with antibody cocktail for surface markers.

### Western blot

Cells were lysed in RIPA Lysis and Extraction buffer (Thermo Fisher Scientific) with 1:100 protease inhibitor cocktail (Sigma Aldrich) on ice for 10 minutes and then centrifuged at max speed for 10 minutes. Supernatants were mixed at 1:1 (vol/vol) with 2X Laemmli Sample Buffer (Biorad) with 0.1M dithiothreitol (DTT) and boiled for 5 minutes. Samples were then separated with sodium dodecyl sulfate polyacrylamide gel electrophoresis (SDS-PAGE) for 1.5 hours at 150V and transferred to nitrocellulose membranes for 1 hour at 95V. Membranes were then blocked with 4% (wt/vol) milk powder and 1% (wt/vol) bovine serum albumin (BSA) in 0.1% Tween 20 (vol/vol) phosphate buffered saline (PBST) for 3 hours. Membranes were then incubated with the listed primary antibodies overnight at 4°C, p57^Kip2^ (BD Biosciences), p27^Kip1^ (BD Biosciences), NR4A1 (BD Biosciences), GAPDH (Millipore) and α-globin (Santa Cruz Biotechnologies). Membranes were washed 5 time for 5 minutes with PBST and incubated with horseradish peroxidase (HRP) conjugated secondary antibodies (Biorad) for 2 hours at room temperature. Membranes were imaged with Pierce ECL Western Blotting Substrate (Thermo Fisher Scientific) using a ChemiDoc MP Imaging System (Biorad). Western blot images are representative of multiple experiments and were quantified with ImageJ (NIH).

### Lentiviral transduction

p57^Kip2^ and p27^Kip1^ knockdown experiments were each carried out with 2 lentiviral shRNA knockdown constructs targeting CDKN1C and CDKN1B respectively (Clone ID: NM_000076.2-1451s21c1 and NM_000076.2-1216s21c1 for CDKN1C and NM_004064.3-841s21c1 and NM_004064.3-643s21c1 for CDKN1B, Sigma Aldrich). shRNA knockdown constructs targeting luciferase were used as controls. After 2 days of culture in serum free expansion media, erythroid progenitors were placed in 10% FBS IMDM with 3U/mL heparin and 8 μg/mL polybrene. Lentiviral particles were then added at a multiplicity of infection (MOI) of 30 and spinoculated for 2 hours at 3000 rpm. Cells were then incubated overnight and placed in serum free expansion media. After 24 hour recovery, lentiviral transduction was positively selected with 1 μg/mL puromycin which was maintained until day 11 of culture.

### Proteomic profiling

#### In-Solution Digestion

CD34+ cell pellets were lysed for 30 min at 4 °C in 8M urea, 50 mM Tris-HCl pH 8.0, 75 mM NaCl, 1 mM EDTA, 2 μg/μl aprotinin (Sigma-Aldrich), 10 μg/μl leupeptin (Roche), and 1 mM phenylmethylsulfonyl fluoride (PMSF) (Sigma-Aldrich). Lysates were cleared via centrifugation at 20,000 rcf, and protein concentration was determined using a bicinchoninic acid (BCA) protein assay (Pierce). Remaining lysis buffer was used to equalize sample concentration to the lowest measured concentration before proceeding. Protein reduction was performed with 5 mM dithiothreitol (DTT) for 1 h at room temperature, followed by alkylation with 10 mM iodoacetamide for 45 min at room temperature in the dark. Sample volumes were then adjusted with 50 mM Tris-HCl pH 8.0 to reduce urea concentration to 2 M preceding enzymatic digestion. Proteins were digested first with endoproteinase LysC (Wako Laboratories) for 2 h at 25 °C, then overnight with sequencing-grade trypsin (Promega) at 25 °C, both at enzyme-to-substrate ratios of 1:50. Following digestion, samples were acidified to a concentration of 1% with neat formic acid, and insoluble peptides and urea was removed via centrifugation at 20,000 rcf. Remaining soluble peptides were desalted using a 100 mg reverse phase tC18 SepPak cartridge (Waters). Cartridges were conditioned with 1 ml 100% MeCN and 1 ml 50% MeCN/0.1% FA, then equilibrated with 4X 1 ml 0.1% TFA. Samples were loaded onto the cartridge and washed 3X with 1 ml 0.1% TFA and 1X with 1 ml 1% FA, then eluted with 2X 600 μl 50% MeCN/0.1% FA. Peptide concentration of desalted samples was again estimated with a BCA assay such that the proper amount for TMT labeling could be removed, dried in a vacuum centrifuge, and stored at −80 °C.

#### TMT labeling of peptides

Samples were divided into two groups— peripheral blood- and cord blood-derived cells— such that each set could be separately labeled with TMT 6-plex isobaric mass tagging reagents (Thermo Fisher Scientific) as previously described (41). Each 6-plex contained triplicate samples of dexamethasone- and DMSO-treated cells. Digested peptides were resuspended in 50 mM HEPES, pH 8.5 at a concentration of 2.5 mg/ml. Dried TMT reagent was reconstituted at 20 μg/μl in 100% anhydrous MeCN and added to samples at a 1:1 TMT to peptide mass ratio (100 μg for peripheral blood samples and 70 μg for cord blood samples due to limiting material amount). Labeling was performed for 1 hr at 25 °C with shaking. The TMT reaction was quenched with 5% hydroxylamine to a final concentration of 0.2%, shaking for 15 min at 25 °C. TMT-labeled samples within each plex were then combined, dried to completion via vacuum centrifugation, reconstituted in 1 ml 0.1% FA and desalted with a 100 mg SepPak cartridge as described above.

#### Basic Reverse Phase (bRP) Fractionation

TMT-labeled peptides were fractionated via offline basic reverse-phase (bRP) chromatography as previously described (Mertins et al., 2018). Chromatography was performed with a Zorbax 300 Extend-C18 column (4.6 x 250 mm, 3.5 μm, Agilent) on an Agilent 1100 high pressure liquid chromatography (HPLC) system. Samples were reconstituted in 900 μl of bRP solvent A (5 mM ammonium formate, pH 10.0 in 2% vol/vol MeCN) and injected with this solvent at a flow rate of 1 ml/min. Peptides were separated at the same flow rate with a 96 min gradient, beginning with an initial increase to 16% bRP solvent B (5 mM ammonium formate, pH 10.0 in 90% vol/vol MeCN) followed by a linear 60 min gradient to 40% and stepwise ramping to 44% and finally 60% bRP solvent B. A total of 96 fractions were collected in a row-wise snaking pattern into a Whatman 2 ml 96-well plate (GE Healthcare), which were then concatenated non-sequentially into a final 24 fractions for proteomic analysis. Fractions were dried via vacuum centrifugation.

#### Liquid chromatography and mass spectrometry

Dried fractions were reconstituted in 3% MeCN/0.1% FA to a peptide concentration of 1 μg/μl and analyzed via coupled nanoflow liquid chromatography and tandem mass spectrometry (LC-MS/MS) using a Proxeon Easy-nLC 1000 (Thermo Fisher Scientific) and a Q-Exactive Plus series mass spectrometer (Thermo Fisher Scientific). A sample load of 1 μg for each fraction was separated on a capillary column (360 μm outer diameter x 75 μm inner diameter) containing an integrated emitter tip and heated to 50 °C and packed to a length of approximately 30 cm with ReproSil-Pur C18-AQ 1.9 μm beads (Dr. Maisch GmbH)). Chromatography was performed with a 110 min gradient consisting of solvent A (3% MeCN/0.1% FA) and solvent B (90% MeCN/0.1% FA). The gradient profile, described as min:% solvent B, was 0:2, 1:6, 85:30, 94:60, 95:90, 100:90, 101:50, 110:50, with the first six steps being performed at a flow rate of 200 nl/min and the last two at a flow rate of 500 nl/min. Ion acquisition on the Q-Exactive Plus was performed in data-dependent mode, acquiring HCD-MS/MS scans at a resolution of 17,500 on the top 12 most abundant precursor ions in each full MS scan (70,000 resolution). The automatic gain control (AGC) target was set to 3 x 10^6^ ions for MS1 and 5 x 10^4^ for MS2 and the maximum ion time was set to 120 ms for MS2. The collision energy was set to 30, peptide matching was set to preferred, isotope exclusion was enabled, and dynamic exclusion time was set to 20 s.

Raw mass spectrometry data will be made publicly available in MassIVE upon acceptance of the manuscript.

#### Data Analysis

Data was analyzed using Spectrum Mill, version 6.01.202 (Agilent Technologies). In extracting spectra from the .raw format for MS/MS searching, spectra from the same precursor, or within a retention time window of +/- 60 s and m/z range of +/- 1.4 were merged. Spectra were filtered to include only those with a precursor mass range of 750 to 6000 Da and a sequence tag length greater than 0. MS/MS searching was performed against a human UniProt database. Digestion enzyme conditions were set to “Trypsin allow P” for the search, allowing up to 4 missed cleavages within a matched peptide. Fixed modifications were carbamidomethylation of cysteine and TMT6 on the N-terminus and internal lysine. Variable modifications were oxidized methionine and acetylation of the protein N-terminus. Matching criteria included a 30% minimum matched peak intensity and a precursor and product mass tolerance of +/- 20 ppm. Peptide-level matches were validated if found to be below the 1.0% false discovery rate (FDR) threshold and within a precursor charge range of 2-6. A second round of validation was then performed for protein-level matches, requiring a minimum protein score of 0. Protein-centric information, including experimental ratios, was then summarized in a table, which was quality filtered for non-human contaminants, keratins, and any proteins not identified by at least two fully quantified peptides with two ratio counts.

### Statistical analysis

All statistical evaluations between the different experimental groups were performed using GraphPad Prism 8 (unpaired 2-tailed Student’s t-test). A P<0.05 was considered as statistically significant.

For proteomics analyses, peripheral and cord blood plexes were analyzed separately, each utilizing biologically paired samples to compare dexamethasone treatment to a DMSO control. Data was median normalized and subjected to a one-sample moderated T-test using an internal R-Shiny package based in the limma R library. Correction for multiple testing was performed using the Benjamini-Hochberg false discovery rate method.

### Human Studies

All human studies have been approved by the Institutional Review Board (Northwell Health and Stanford University). CD34^+^ cells were obtained from deidentified control adult peripheral blood leukoreduction filters, deidentified cord blood units, or phlebotomized patients with DBA after informed consent was obtained in writing and prior inclusion in the study. In order to limit sample variability, blood from multiple control peripheral blood or cord blood donors were pooled.

Patients with DBA were defined as transfusion-dependent or steroid-responsive based on their clinical need for chronic red blood cells transfusion or successful management with corticosteroids.

The three patients with DBA presented in Figure 1E were patients followed after diagnosis and initial treatment with prednisone. The dose of prednisone was decreased after initial response.

## Author Contributions

R.A. and H.Y. designed, performed most of the experiments, analyzed data and contributed to the writing of the manuscript. N.W., J.H., B.M.D., J.P. performed experiments. M.O., N.D.U., S.A.C., designed, performed, analyzed the proteomics studies and edited the manuscript. A.V., J.M.L., L.D.C., analyzed data related to patients with Diamond Blackfan anemia. C.H., S.K., N.T. designed, analyzed data and edited the manuscript. N.M., A.N. and L.B. designed the project, analyzed the data, and wrote the manuscript.

## Acknowledgements

The authors thank the patients and their families for their contributions. We thank Philippe Marambaud for helpful discussions and critical reading of the manuscript.

This research was supported in part by National Institutes of Health grants DK26263 and DK32094 (to N.M.), HL079571 (to J.M.L.), CA210986, CA214125 (to S.A.C.), CA210979 (to D.R.M.) and HL144436 (to L.B. and A.N.), and by the Pediatric Cancer Foundation (to J.M.L. and L.B.). A.N. is the recipient of an ASH Bridge award and a Faculty Scholar grant from the Maternal and Child Health Research Institute at Stanford University. L.B. is the recipient of a St. Baldrick’s Scholar award.

---

**Supplemental Figure 1:**
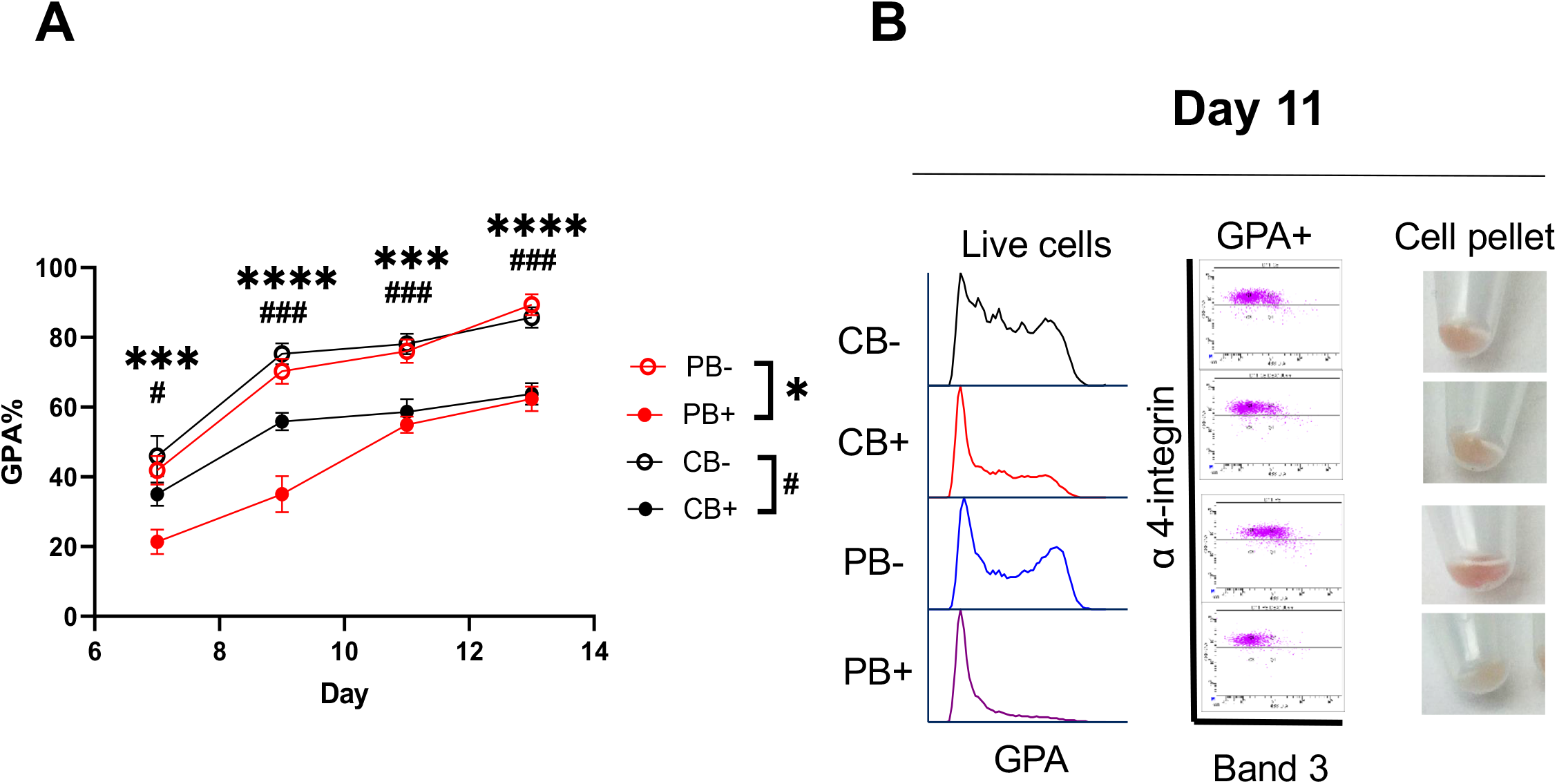
(A) Quantification of the percentage of GPA positive cells in PB (n=4) or CB (n=4) cultures under expansion with or without Dex. PB is shown in red while CB is shown in black. Dex treated cultures are shown as solid while untreated cultures as shown as open. (B) Representative flow cytograms and cell pellets of PB or CB cultures with or without Dex treatment at day 11 of differentiation. PB and CB-derived cells have slower accumulation of GPA and Band 3 expression. Packed PB-derived cells treated with Dex appear paler due to delayed production of hemoglobin. Data for B represent 1 of 3 independent experiments. Data for A are shown as mean ± SE. *P < 0.05, ***P < 0.001, ****P < 0.0001, by 2-tailed Student’s *t* test with Holm-Sidak correction for multiple comparisons (A).

**Supplemental Figure 2:**
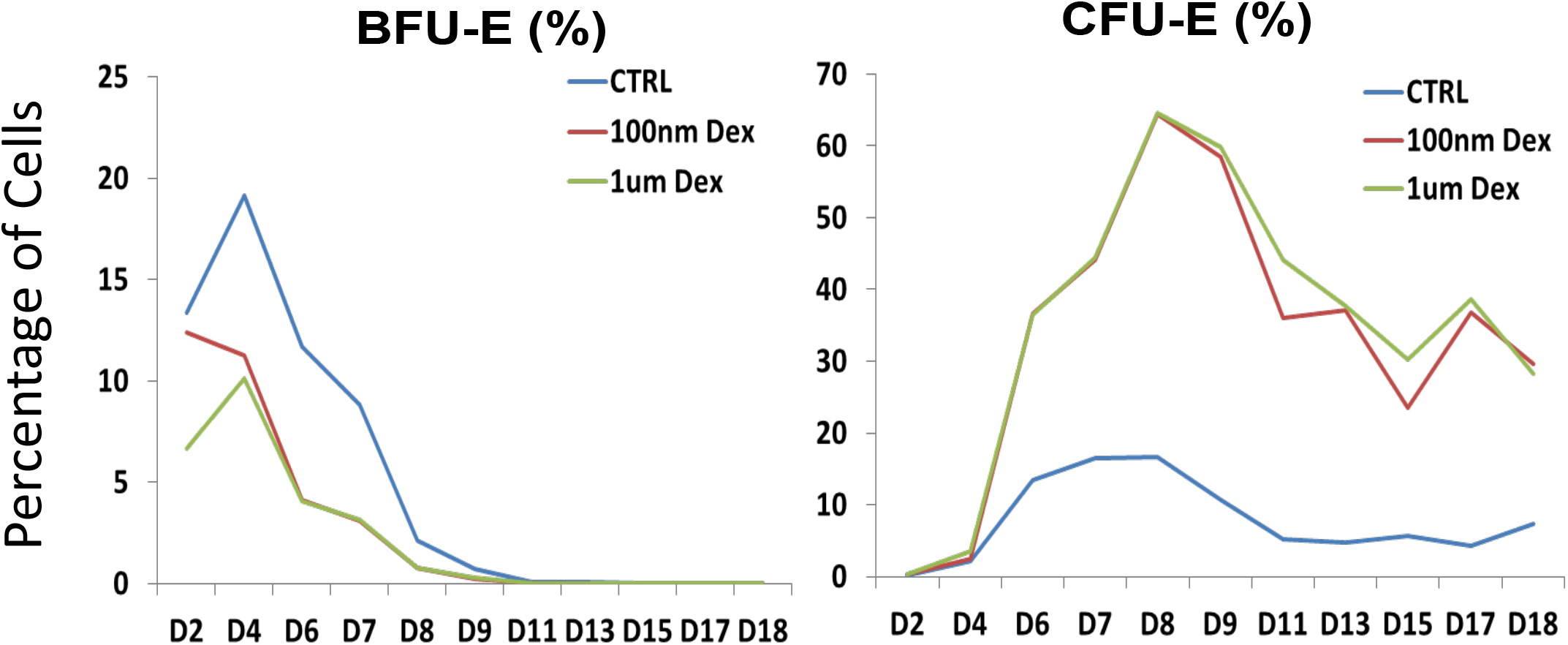
Percentage of BFU-E and CFU-E from PB-derived CD34+ cultures during expansion with varying concentrations of Dex. No additional effect is seen from concentrations of Dex greater than 100 nM.

**Supplemental Figure 3:**
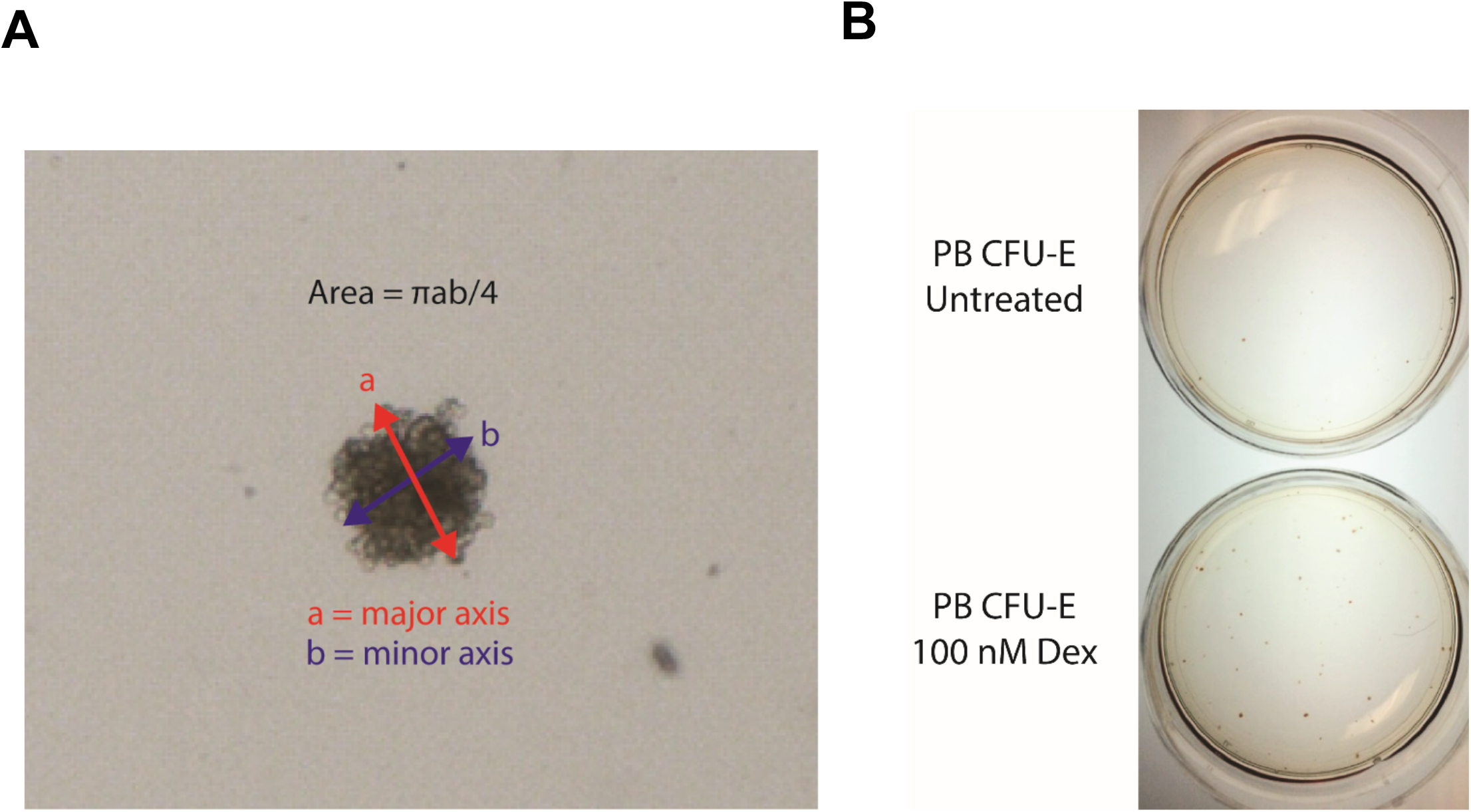
(A) Example of CFU-E colony area measurement. Colony morphology was modeled as an ellipse and the area was determined by measuring its major axis a and minor axis b to calculate area by the formula A = πab/4. (B) Representative image of CFU-E colonies formed from sorted PB derived CFU-E with or without Dex treatment.

**Supplemental Figure 4:**
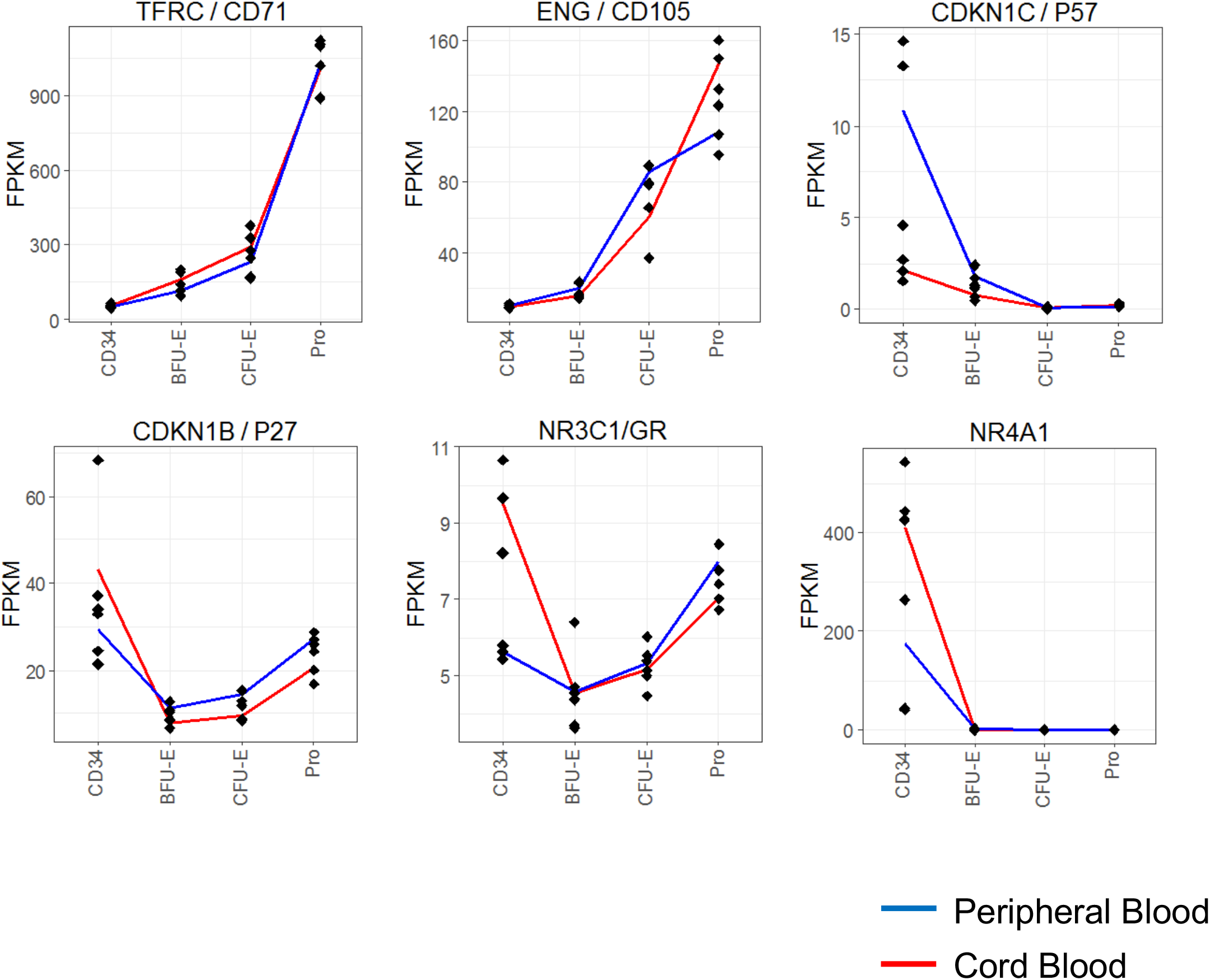
Quantification of RNA expression of TFRC (CD71), ENG (CD105), CDKN1C (p57^Kip2^), CDKN1B (p27^Kip1^), NR3C1 (GR), and NR4A1 in sorted erythroid populations. PB is shown in blue while CB is shown in red.

**Supplemental Figure 5:**
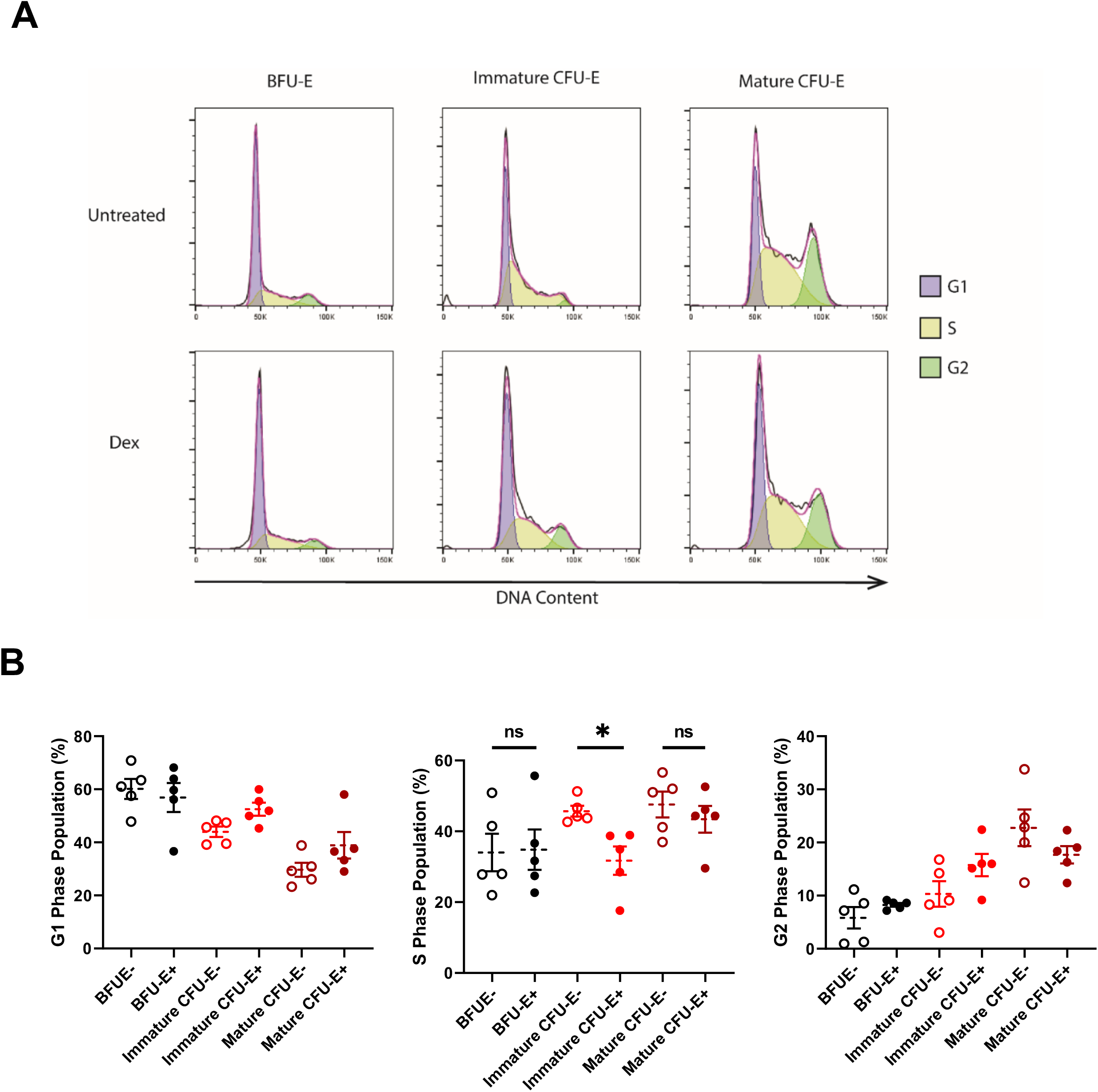
(A) Representative cell cycle profiles of BFU-E, immature CFU-E and mature CFU-E derived from PB with or without Dex treatment and stained with Hoechst 33342. G1 is shown in purple, S is shown yellow and G2 is shown in green. (B) Quantification of the population of cells in G1, S, and G2 phases for BFU-E, immature CFU-E and mature CFU-E derived from PB treated with or without Dex and stained with Hoechst 33342 (n = 5). Data for A represents 1 of 5 independent experiments. Data for B are shown as mean ± SE. *P < 0.05, by 2-tailed Student’s *t* test (B).

**Supplemental Figure 6:**
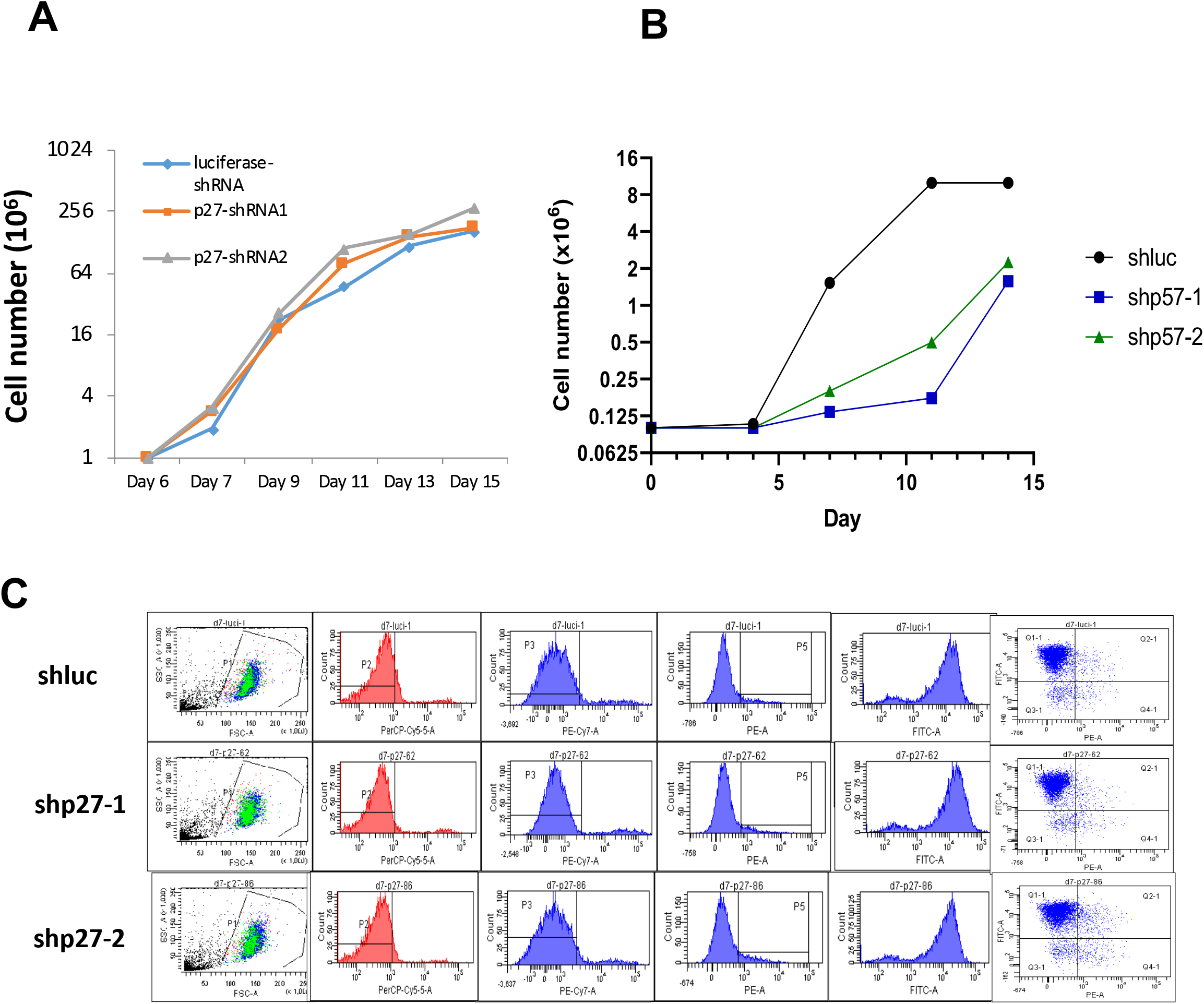
(A) Growth curves of PB CD34^+^ cells transduced with lentivirus for shRNA knockdown of p27^Kip1^ and luciferase control. (B) Growth curves of PB CD34^+^ cells transduced with lentivirus for shRNA knockdown of p57^Kip2^ and luciferase control with or without Dex treatment. (C) Flow cytograms of PB CD34^+^ cells transduced with lentivirus for shRNA knockdown of p27^Kip1^ and luciferase control. Data for A, B and C represent 1 of 3 independent experiments.

**Supplemental Table 1:**

List of proteins identified by proteomics in peripheral blood and cord blood.

## References

1. Da Costa L, Moniz H, Simansour M, Tchernia G, Mohandas N, and Leblanc T. Diamond-Blackfan anemia, ribosome and erythropoiesis. Transfus Clin Biol. 2010;17(3):112–9.

2. Vlachos A, and Muir E. How I treat Diamond-Blackfan anemia. Blood. 2010;116(19):3715–23.

3. Chan HS, Saunders EF, and Freedman MH. Diamond-Blackfan syndrome. II. In vitro corticosteroid effect on erythropoiesis. Pediatr Res. 1982;16(6):477–8.

4. Kadmiel M, and Cidlowski JA. Glucocorticoid receptor signaling in health and disease. Trends Pharmacol Sci. 2013;34(9):518–30.

5. Danilova N, and Gazda HT. Ribosomopathies: how a common root can cause a tree of pathologies. Dis Model Mech. 2015;8(9):1013–26.

6. Vlachos A, Blanc L, and Lipton JM. Diamond Blackfan anemia: a model for the translational approach to understanding human disease. Expert Rev Hematol. 2014;7(3):359–72.

7. Khajuria RK, Munschauer M, Ulirsch JC, Fiorini C, Ludwig LS, McFarland SK, et al. Ribosome Levels Selectively Regulate Translation and Lineage Commitment in Human Hematopoiesis. Cell. 2018;173(1):90–103 e19.

8. Ludwig LS, Gazda HT, Eng JC, Eichhorn SW, Thiru P, Ghazvinian R, et al. Altered translation of GATA1 in Diamond-Blackfan anemia. Nat Med. 2014;20(7):748–53.

9. Quarello P, Garelli E, Carando A, Brusco A, Calabrese R, Dufour C, et al. Diamond-Blackfan anemia: genotype-phenotype correlations in Italian patients with RPL5 and RPL11 mutations. Haematologica. 2010;95(2):206–13.

10. Hwang Y, Futran M, Hidalgo D, Pop R, Iyer DR, Scully R, et al. Global increase in replication fork speed during a p57KIP2-regulated erythroid cell fate switch. Sci Adv. 2017;3(5):e1700298.

11. Lee HY, Gao X, Barrasa MI, Li H, Elmes RR, Peters LL, et al. PPAR-alpha and glucocorticoid receptor synergize to promote erythroid progenitor self-renewal. Nature. 2015;522(7557):474–7.

12. Li H, Natarajan A, Ezike J, Barrasa MI, Le Y, Feder ZA, et al. Rate of Progression through a Continuum of Transit-Amplifying Progenitor Cell States Regulates Blood Cell Production. Dev Cell. 2019;49(1):118–29 e7.

13. Zhang L, Prak L, Rayon-Estrada V, Thiru P, Flygare J, Lim B, et al. ZFP36L2 is required for self-renewal of early burst-forming unit erythroid progenitors. Nature. 2013;499(7456):92–6.

14. Gregory CJ, and Eaves AC. Human marrow cells capable of erythropoietic differentiation in vitro: definition of three erythroid colony responses. Blood. 1977;49(6):855–64.

15. Gregory CJ, and Eaves AC. Three stages of erythropoietic progenitor cell differentiation distinguished by a number of physical and biologic properties. Blood. 1978;51(3):527–37.

16. McLeod DL, Shreeve MM, and Axelrad AA. Improved plasma culture system for production of erythrocytic colonies in vitro: quantitative assay method for CFU-E. Blood. 1974;44(4):517–34.

17. Stephenson JR, Axelrad AA, McLeod DL, and Shreeve MM. Induction of colonies of hemoglobin-synthesizing cells by erythropoietin in vitro. Proc Natl Acad Sci U S A. 1971;68(7):1542–6.

18. Li J, Hale J, Bhagia P, Xue F, Chen L, Jaffray J, et al. Isolation and transcriptome analyses of human erythroid progenitors: BFU-E and CFU-E. Blood. 2014;124(24):3636–45.

19. Iskander D, Psaila B, Gerrard G, Chaidos A, En Foong H, Harrington Y, et al. Elucidation of the EP defect in Diamond-Blackfan anemia by characterization and prospective isolation of human EPs. Blood. 2015;125(16):2553–7.

20. Flygare J, Rayon Estrada V, Shin C, Gupta S, and Lodish HF. HIF1alpha synergizes with glucocorticoids to promote BFU-E progenitor self-renewal. Blood. 2011;117(12):3435–44.

21. Lodish H, Flygare J, and Chou S. From stem cell to erythroblast: regulation of red cell production at multiple levels by multiple hormones. IUBMB Life. 2010;62(7):492–6.

22. Gao X, Lee HY, da Rocha EL, Zhang C, Lu YF, Li D, et al. TGF-beta inhibitors stimulate red blood cell production by enhancing self-renewal of BFU-E erythroid progenitors. Blood. 2016;128(23):2637–41.

23. Golde DW, Bersch N, and Cline MJ. Potentiation of erythropoiesis in vitro by dexamethasone. J Clin Invest. 1976;57(1):57–62.

24. Ohene-Abuakwa Y, Orfali KA, Marius C, and Ball SE. Two-phase culture in Diamond Blackfan anemia: localization of erythroid defect. Blood. 2005;105(2):838–46.

25. Dulmovits BM, Appiah-Kubi AO, Papoin J, Hale J, He M, Al-Abed Y, et al. Pomalidomide reverses γ-globin silencing through the transcriptional reprogramming of adult hematopoietic progenitors. Blood. 2016;127(11):1481–92.

26. Nathan DG, and Gardner FH. Erythroid cell maturation and hemoglobin synthesis in megaloblastic anemia. J Clin Invest. 1962;41:1086–93.

27. Davies SV, Cavill I, Bentley N, Fegan CD, Poynton CH, and Whittaker JA. Evaluation of erythropoiesis after bone marrow transplantation: quantitative reticulocyte counting. Br J Haematol. 1992;81(1):12–7.

28. d’Onofrio G, Chirillo R, Zini G, Caenaro G, Tommasi M, and Micciulli G. Simultaneous measurement of reticulocyte and red blood cell indices in healthy subjects and patients with microcytic and macrocytic anemia. Blood. 1995;85(3):818–23.

29. Yan H, Hale J, Jaffray J, Li J, Wang Y, Huang Y, et al. Developmental differences between neonatal and adult human erythropoiesis. Am J Hematol. 2018;93(4):494–503.

30. Matsumoto A, Takeishi S, Kanie T, Susaki E, Onoyama I, Tateishi Y, et al. p57 is required for quiescence and maintenance of adult hematopoietic stem cells. Cell Stem Cell. 2011;9(3):262–71.

31. Hsieh FF, Barnett LA, Green WF, Freedman K, Matushansky I, Skoultchi AI, et al. Cell cycle exit during terminal erythroid differentiation is associated with accumulation of p27(Kip1) and inactivation of cdk2 kinase. Blood. 2000;96(8):2746–54.

32. Reddy TE, Gertz J, Crawford GE, Garabedian MJ, and Myers RM. The hypersensitive glucocorticoid response specifically regulates period 1 and expression of circadian genes. Mol Cell Biol. 2012;32(18):3756–67.

33. Freire PR, and Conneely OM. NR4A1 and NR4A3 restrict HSC proliferation via reciprocal regulation of C/EBPα and inflammatory signaling. Blood. 2018;131(10):1081–93.

34. D’Arena G, Musto P, Cascavilla N, Di Giorgio G, Zendoli F, and Carotenuto M. Human umbilical cord blood: immunophenotypic heterogeneity of CD34+ hematopoietic progenitor cells. Haematologica. 1996;81(5):404–9.

35. Fauser AA, and Messner HA. Fetal hemoglobin in mixed hemopoietic colonies (CFU-GEMM), erythroid bursts (BFU-E) and erythroid colonies (CFU-E): assessment by radioimmune assay and immunofluorescence. Blood. 1979;54(6):1384–94.

36. Nathan DG, Chess L, Hillman DG, Clarke B, Breard J, Merler E, et al. Human erythroid burst-forming unit: T-cell requirement for proliferation in vitro. J Exp Med. 1978;147(2):324–39.

37. Tusi BK, Wolock SL, Weinreb C, Hwang Y, Hidalgo D, Zilionis R, et al. Population snapshots predict early haematopoietic and erythroid hierarchies. Nature. 2018;555(7694):54–60.

38. Pauklin S, and Vallier L. The cell-cycle state of stem cells determines cell fate propensity. Cell. 2013;155(1):135–47.

39. Palumbo-Zerr K, Zerr P, Distler A, Fliehr J, Mancuso R, Huang J, et al. Orphan nuclear receptor NR4A1 regulates transforming growth factor-β signaling and fibrosis. Nat Med. 2015;21(2):150–8.

40. Ge J, Apicella M, Mills JA, Garçon L, French DL, Weiss MJ, et al. Dysregulation of the Transforming Growth Factor β Pathway in Induced Pluripotent Stem Cells Generated from Patients with Diamond Blackfan Anemia. PLoS One. 2015;10(8):e0134878.

41. Zecha J, Satpathy S, Kanashova T, Avanessian SC, Kane MH, Clauser KR, et al. TMT Labeling for the Masses: A Robust and Cost-efficient, In-solution Labeling Approach. Mol Cell Proteomics. 2019;18(7):1468–78.

